# sgRNA level is a major factor affecting CRISPRi knockdown efficiency in K562 cells

**DOI:** 10.1101/2020.01.12.903625

**Authors:** Y. Wang, Y. Xie, Z. C. Dong, X. J. Jiang, P. Gong, J. Lu, F. Wan

**Affiliations:** College of Life Sciences, Inner Mongolia Agricultural University, Inner Mongolia, 010010 China; College of Science, Inner Mongolia Agricultural University, Inner Mongolia, 010010 China

**Keywords:** CRISPR interference, knockdown efficiency, inducible Tet-on system, the multiplicity of infection, sgRNA expression level

## Abstract

To determine how nuclease deactivated Cas9 (dCas9) or sgRNA expression level affects the knockdown efficiency of CRISPRi, K562 cell clones expressing KRAB-dCas9 protein either with the inducible Tet-on system or with the constitutive SFFV promotor were created by lentiviral transduction, and single clones were selected by fluorescence-activated cell sorting (FACS) for further study. Six genes with various expression levels were targeted using lentiviral sgRNA from two libraries in four cell clones with various KRAB-dCas9 expression levels. We determined the knockdown efficiency and the expression level of the dCas9 protein /sgRNA level by flow cytometry. The cell clone with the highest KRAB-dCas9 expression level achieved effective CRISPRi knockdown, and is statistically different from other clones, indicating enough KRAB-dCas9 expression might be a prerequisite for CRISPRi. Utilizing this clone, we modified the expression level of sgRNA by adopting different multiplicity of infection (MOI)in lentiviral transduction and found that the knockdown efficiency was neither affected by the target gene expression level nor does it correlate with KRAB-dCas9 level, which remained relatively constant (CV=2.2%) across knockdown experiments. 74.72%, 72.28%, 39.08% knockdown of *mmadhc, rpia, znf148* genes were achieved, and the knockdown efficiency correlated well with the sgRNA expression level. Linear regression modeling of the data revealed that the knockdown efficiency is significantly affected by both KRAB-dCas9 and sgRNA level, and the sgRNA level has a greater impact, based on the standardized coefficient (0.525 for KRAB-dCas9, 0.981 for sgRNA), indicating that sgRNA level is a major factor affecting CRISPRi efficiency.

## INTRODUCTION

CRISPRi is gradually replacing the siRNA technique for mechanistic investigations in various fields [1]. However, there is a lack of mechanistic study on the efficiency of the CRISPRi system. How the two major players of the system, the dCas9 fusion protein, and the guide RNA affect the knockdown efficiency has not been systematically studied.

Three protein domains were commonly used for repressing target gene transcription: KRAB, MeCP2, dCas9 and its synthetic derivatives [2-3]. KRAB recruits KAP1, which serves as a scaffold for various heterochromatin-inducing factors [4]; MeCP2 binds to the methylated DNA and interacts with histone deacetylase and the co-repressor SIN3A [5]. A combination of dCas9 with KRAB and MeCP2 has been shown to improve the knockdown efficiency [2-3], and dCas9, KRAB-dCas9, dCas9-KRAB-MeCP2 are three commonly used combinations.

Although the dynamics of KRAB-dCas9 and sgRNA mediated CRISPRi knockdown has not been studied thoroughly, the mechanistic study of the CRISPR system could provide some clues due to the similarity: effective knockdown of target gene started from the sgRNA being expressed from the transgene, formed a complex with Cas9 protein, and roaming in the nucleus until the sgRNA hybrid to target DNA, then the Cas9 protein cut the target gene [6]. Cas9 is essential for guide RNA stability and that the nuclear Cas9–guide RNA complex levels limit the targeting efficiency. Similar dynamics could exist in the CRISPRi system, and the very first step is to determine how dCas9 fusion protein and the sgRNA expression level affect CRISPRi efficiency.

In this study, different cell clones with various KRAB-dCas9 protein level and subclones with different sgRNA expression level was evaluated for CRISPRi efficiency. Linear regression was applied to determine the major factor that determines CRISPRi efficiency.

## EXPERIMENTAL

### Cell culture and construction of plasmids

HEK293T and K562 cells were obtained from the National Infrastructure of Cell Line Resource. HEK293T cells were cultured at 37°C in Dulbecco’s modified Eagle’s medium (DMEM) containing 4.5 g/L glucose (Gibco) supplemented with 10% (v/v) fetal bovine serum (FBS, Gibco). K562 cells were maintained in Iscove’s modified Dulbecco’s medium (IMDM, Gibco) supplemented with 10% FBS. Cells were tested every 3 months for mycoplasma contamination and consistently tested negative. To induce the expression of KRAB-dCas9 protein using doxycycline, we replaced the promotor of SFFV-KRAB-dCas9 plasmid (plasmid Addgene, #60954 provided by Jonathan S. Weissman) with the pTRE3G promoter (plasmid Addgene # 96964 provided by Elena Cattaneo) using classic cloning method, primers are listed in Supplementary Table 1.

**Table 1.**
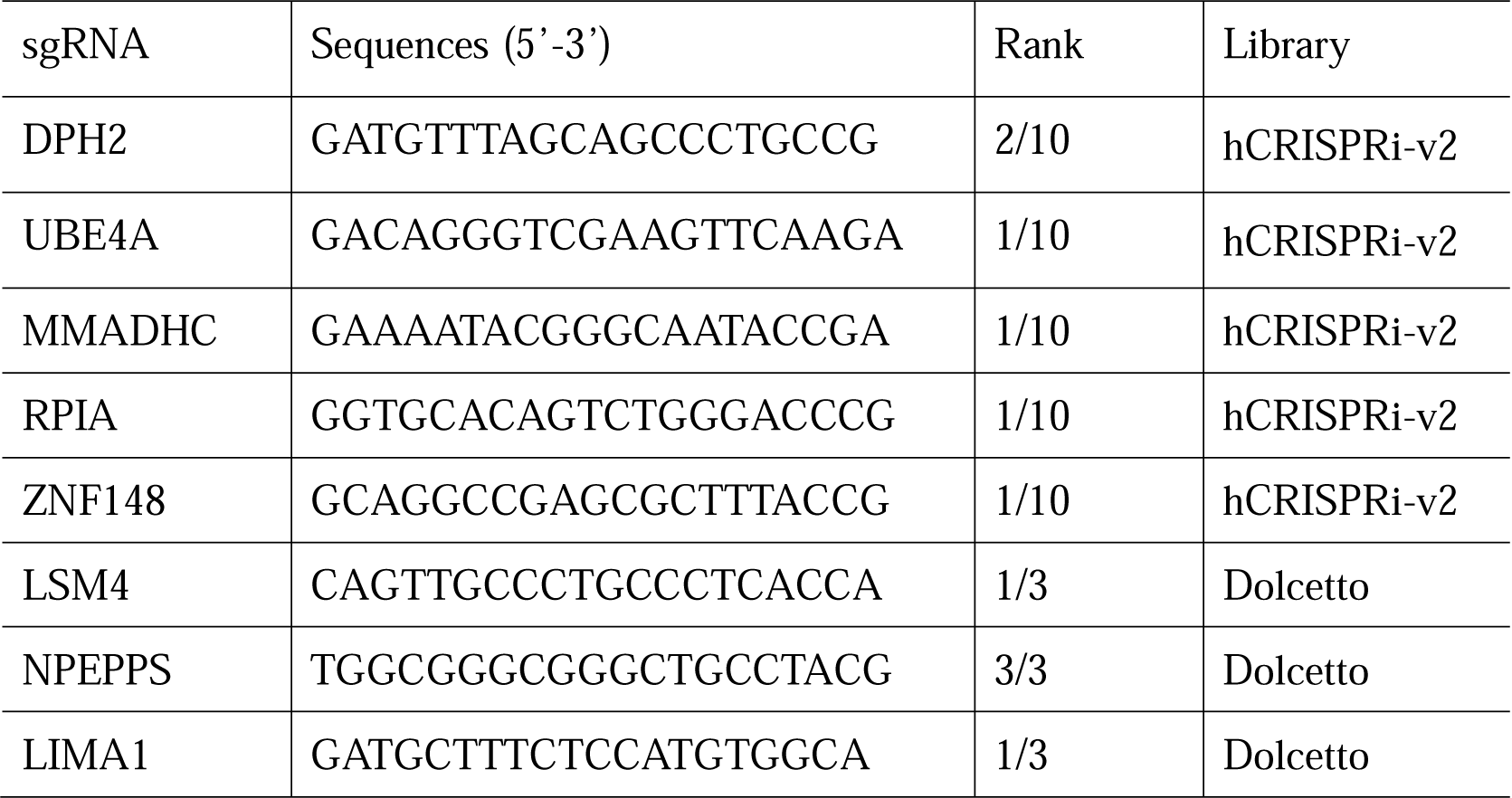
List of sgRNA sequences

### Lentiviral Packaging

2.1×10^7^ HEK293T cells were seeded into T175 flasks 1d before transfection. Cells were transfected with the packaging plasmids of pSPAX2 (11.6 ug) and pMD2.G (2.8 ug) (plasmid Addgene #12260 and #12259 provided by Didier Trono), and the target plasmids (11.6 ug) using Polyetherimide (PEI) (1 mg/mL), with the nitrogen to phosphorus (N/P) ratio 20 [2, 7]. Viral supernatants were collected at 48 h and 72 h after transfection and filtered using 0.45 um filters. Subsequently, 4 mL 20% sucrose solution was added to the bottom of the 32 mL viral supernatants, and centrifuge for 2 h at 82,700 g, 4°C [8], and the pellet was resuspended by 100 uL pre-cooled PBS without Ca^2+^/Mg^2+^. The titer of the lentivirus was determined using flow cytometry method as described [8].

### Generating cell clones by lentiviral transduction

To generate single clones with constitutive KRAB-dCas9 protein expression, 1×10^5^ K562 cells were transduced with the lentiviral particles for SFFV-KRAB-dCas9 using polybrene (final concentration 10 ug/mL) at MOI 20 in six-well plates. 72 h after transduction, cells were analyzed and sorted into 96-well plates using a cell sorte(r BD FACS Aria II). Single clones emerged after 14 days and were expanded and cultured for further analysis.

For inducible KRAB-dCas9 expressing cells,1×10^5^ K562 cells were infected with lentiviral particles for expressing the trans-activator of the pTRE3G promotor (pLVX-Tet3G) and selected for 10 days using G418 (Final concentration 100 ug/mL). Subsequently, the cell clones were transduced with the lentiviral particles for pTRE3G-KRAB-dCas9 using polybrene, the following step with the same procedure described in the above section.

### Flow cytometry analysis

The flow cytometry analysis was performed on Beckman CytoFLEX S. The data were analyzed using Flowjo 10 and CytoExpert 2.0 software.

### Western blot analysis and ELISA

To detect the expression level of KRAB-dCas9 protein, K562-idCas9 #49 were induced by the doxycycline of 200 nM and 2000 nM for 48 h. Then, total protein was extracted using RIPA lysis buffer with 1 mM PMSF, and concentration was determined by the BCA protein assay kit (Solarbio, PC0020). For Western blot analysis, total protein (20 ug) were separated by 8% SDS-PAGE and transferred onto a 0.22 um nitrocellulose membrane at 150 V, 90 min. The membrane was incubated with the Cas9 polyclonal antibody (Clontech, 632607) and α-Tubulin (ABclonal, AC012), and the secondary antibodies IRDye™-800CW(BD, Lot No. C50113-06). The membrane was scanned by the Odyssey™CLx Imaging system (Li-COR Bioscience). For ELISA, the KRAB-dCas9 protein was quantified using the CRISPR/Cas9 assay ELISA kit (Epigentek, P-4060-48) according to the manufacturer’s protocol.

### The evaluation of the CRISPRi knockdown efficiency

Single clones were infected with the lentivirus of sgRNA (MOI 20 if not explicitly stated otherwise) with polybrene (Final concentration 10 ug/mL). After 72 h transduction, clones were selected by puromycin (Final concentration 5 ug/mL) for 72 h. The expression level of target genes was detected by qPCR using the relative quantification method. qPCR primers are listed in Supplementary Table 1.

### Statistical analysis

ANOVA and multiple comparisons were performed to compare the four-cell clones. Linear regression modeling was performed to analyze the relationship between the knockdown efficiency and the expression level of either KRAB-dCas9 or sgRNA. All statistical analysis was performed using the SPSS package.

## RESULTS

To analyze how KRAB-dCas9 protein and/or the sgRNA expression level affects the CRISPRi knockdown efficiency, we adopted a hierarchical design that firstly compared cell clones with various KRAB-dCas9 expression level, then compared cell subclones with various sgRNA expression level (Fig. 1a). For the first comparison, we established K562 cell clones that have either constitutive or inducible expression of the KRAB-dCas9-mCherry fusion protein.

**Fig. 1.**
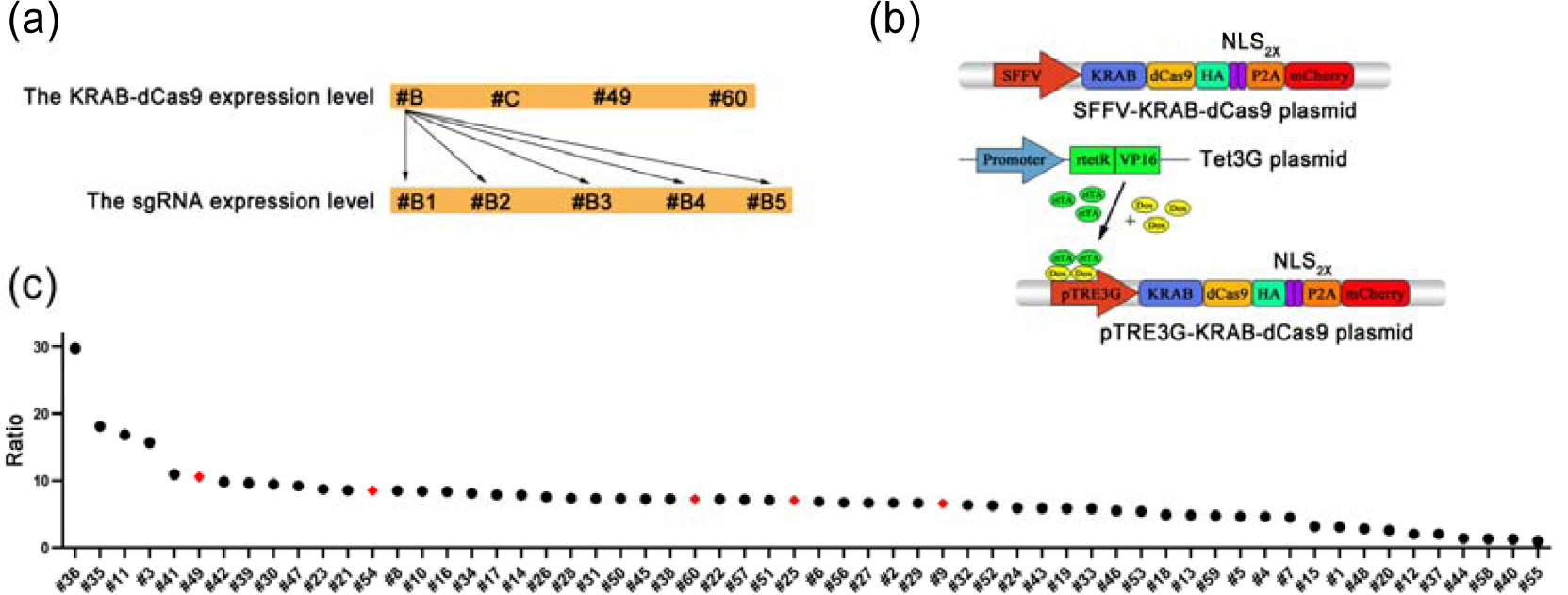
The construction and selection of inducible cell clones. (a) Experiment design. #B: K562-dCas9 #B; #C: K562-dCas9 #C; #49: K562-idCas9 #49; #60: K562-idCas9 #60. #B1, #B2, #B3, #B4, #B5 indicates K562-dCas9 #B infected with difference MOI of sgRNA. (b) Lentiviral vectors for inducible and constitutive KRAB-dCas9 expression. (c) Sixty inducible cell clones ranked based on induced MFI/uninduced MFI, the red prismatic indicated the single clones that chosen for further analysis. 2000 nM doxycycline was applied for induction for 48 h.

Tet-on system was adopted to establish cell clones that express KRAB-dCas9-mCherry fusion protein in doxycycline-inducible fashion: trans-activator (Tet3G plasmid) of the pTRE3G promotor was first introduced into K562 cells by lentiviral transduction, followed by G418 selection, and subsequently undergone lentiviral transduction of the KRAB-dCas9-mCherry expressing vector with the inducible pTRE3G promotor (Fig. 1b). After FACS selection based on mCherry expression, sixty inducible clones (K562-idCas9 #1 to #60) were randomly chosen from twelve 96-well plates for further expansion, and their transgene expression pattern was analyzed with flow cytometry upon doxycycline induction (Supplemental Fig. 1-2). Based on the fold change of transgene expression level, represented by the medium fluorescent intensity (MFI), with or without doxycycline induction (Induced MFI/Uninduced MFI), the 60 clones were ranked (Fig. 1c) and five clones (K562-idCas9 #9, #25, #49, #54, #60) with various fold changes were chosen for further analysis.

We first confirmed the inducible expression of KRAB-dCas9-mCherry in K562-idCas9 #49 and #60 by detecting mCherry, the tag that co-expressed with KRAB-dCas9, using flow cytometry (Fig. 2a), then by detecting dCas9 with western blot and ELISA (Fig. 2b-c). KRAB-dCas9 protein mainly localized in the nucleus, as expected. MFI of mCherry exhibits a similar pattern as the ELISA signal, confirming that mCherry represented the KRAB-dCas9 expression level. For sgRNA, two sgRNA libraries, the hCRISPRi-v2 library [9] and the Dolcetto library [10], were obtained from Addgene, both contain multiple sgRNAs targeting one gene and ranked them (Table 1). Pilot CRISPRi knockdown experiments were carried out using the lentiviral particle of highest-ranking sgRNA against UBE4A and DPH2, from the hCRISPRi-v2 library. K562-idCas9 #9, #25, #54 were excluded from further analysis based on their performance in the pilot experiments (Supplementary Fig. 3a-d). K562-idCas9 #49 displayed 24% knockdown efficiency in the DPH2 gene with 100 nM doxycycline (Supplementary Fig. 3e), has relatively higher KRAB-dCas9-mCherry expression level, and were chosen for further analysis, together with K562-idCas9 #60, which has relatively low KRAB-dCas9-mCherry expression (Fig. 2d).

**Fig. 2.**
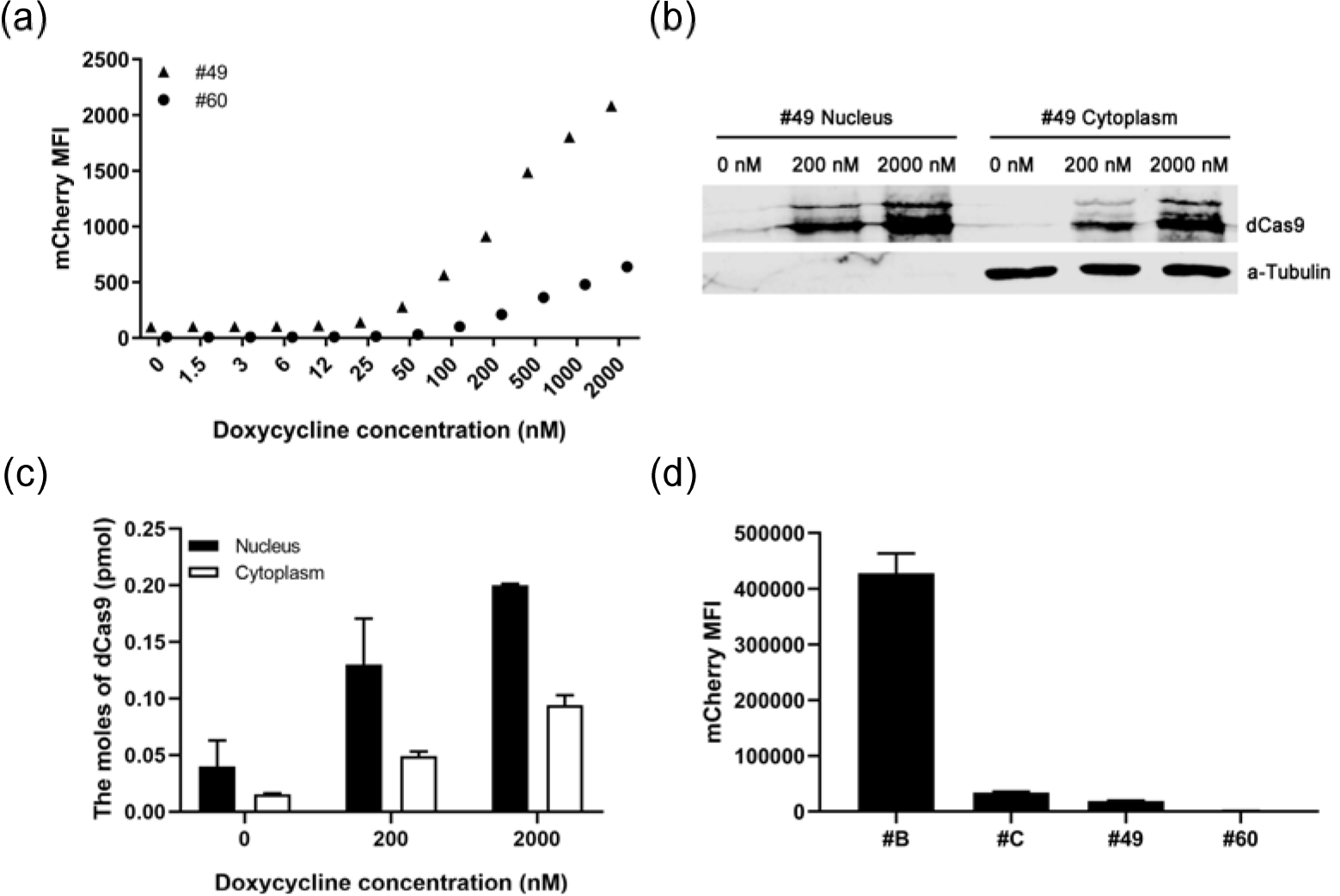
The expression of KRAB-dCas9 in inducible and constitutive cell clones. (a) mCherry MFI in K562-idCas9 #49, K562-idCas9 #60, in response to various doses of doxycycline. Triangle indicated K562-idCas9 #49, round indicated K562-idCas9 #60. (b-c) Western blot and ELISA detection of nuclear and cytoplasmic KRAB-dCas9 in K562-idCas9 #49 induced by 200 nM and 2000 nM doxycycline. (d) mCherry MFI of two inducible cell clones induced by 100 nM doxycycline and two constitutive cell clones. #B: K562-dCas9 #B; #C: K562-dCas9 #C; #49: K562-idCas9 #49; #60: K562-idCas9 #60.

**Fig. 3.**
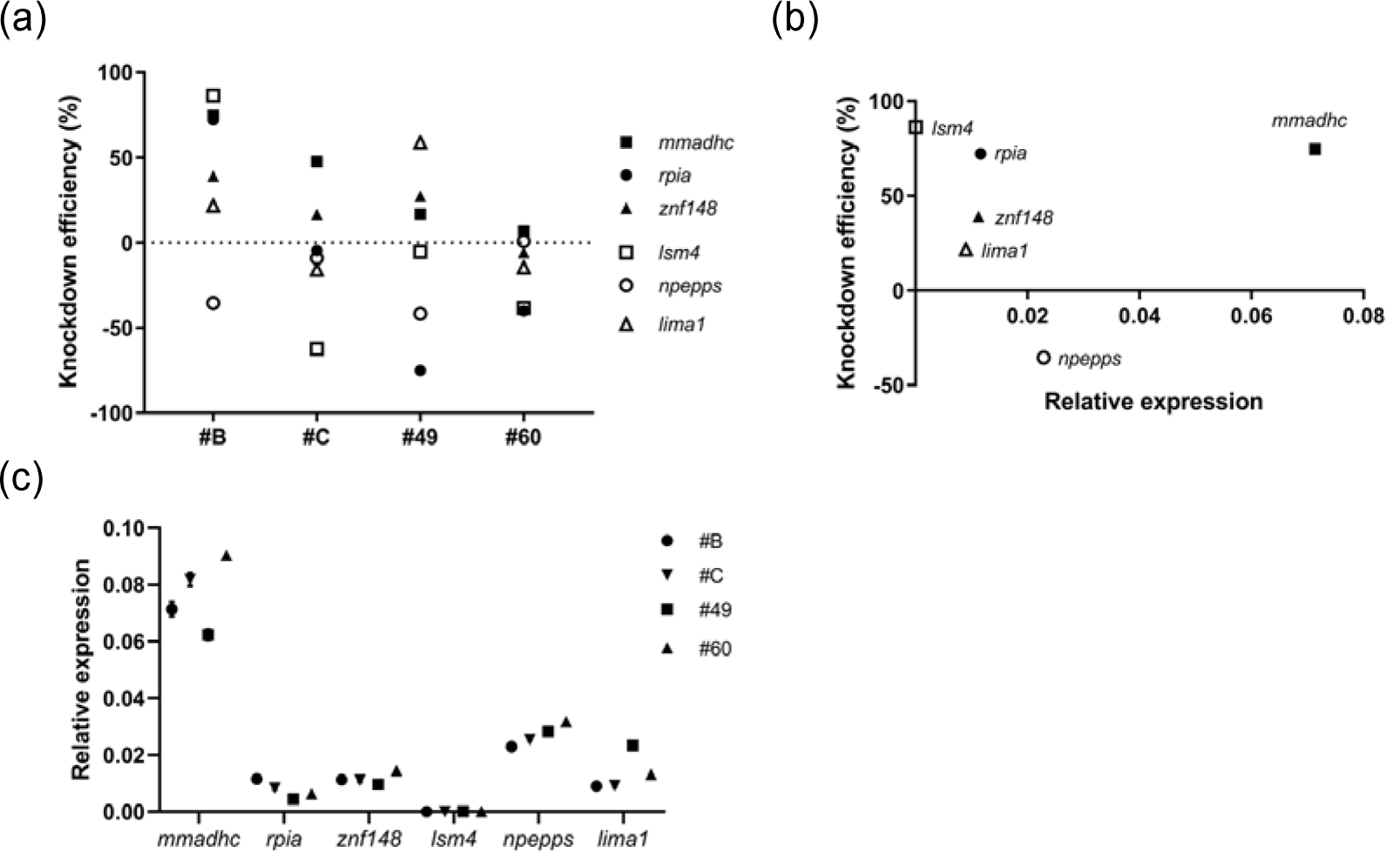
Efficient CRISPRi knockdown requires enough KRAB-dCas9 expression. (a) Knockdown efficiency of six genes in four clones, #B: K562-dCas9 #B; #C: K562-dCas9 #C; #49: K562-idCas9 #49; #60: K562-idCas9 #60. (b) The relationship between target gene expression level and CRISPRi knockdown efficiency in K562-dCas9 #B. (c) The relative expression level of six genes in four clones, #B: K562-dCas9 #B; #C: K562-dCas9 #C; #49: K562-idCas9 #49; #60: K562-idCas9 #60.

Similarly, we established constitutive KRAB-dCas9-mCherry expressing cell clones, and chose three clones, K562-dCas9 #A, K562-dCas9 #B, and K562-dCas9 #C. Pilot knockdown experiments revealed that K562-dCas9 #A failed to knockdown UBE4A or DPH2 genes (Supplemental Fig. 3f), and was excluded from further analysis.

To compare the knockdown efficiency of cell clones with various KRAB-dCas9 expression levels, the constitutive K562-dCas9 #B, #C and the inducible K562-idCas9 #49, #60 were evaluated for their CRISPRi efficiency. The inducible KRAB-dCas9 expressing clones have lower expression levels compared to the constitutive clones, indicating that inducible expression may not be able to reach the expression level of constitutive expression (Fig. 2d). For sgRNA, we randomly choose three sgRNAs per CRISPRi library of high rankings (Table 1). After lentiviral transduction of sgRNA into the cell clones, puromycin selection was applied, and the knockdown efficiency was assessed by qPCR. *mmadhc, rpia, znf148, lsm4*, and *lima1* genes have been knocked down in clone K562-dCas9 #B, the clone with the highest KRAB-dCas9 expression level (Fig. 3a); yet all other clones have relatively low or no knockdown efficiency or the target gene expression level increased (Fig. 3a), indicating sufficient KRAB-dCas9 expression level is a prerequisite for efficient CRISPRi.

To better understanding the difference among the four cell clones, ANOVA was performed and multiple comparisons were made. The results indicated that K562-dCas9 #B is significantly different from K562-dCas9 #C and K562-idCas9 #60, and the p-value for comparison between K562-dCas9 #B and K562-idCas9 #49 is 0.054, indicating that K562-dCas9 #B is also different from K562-idCas9 #49, corroborating our finding that only K562-dCas9 #B has effective CRISPRi (Table 2).

**Table 2.**
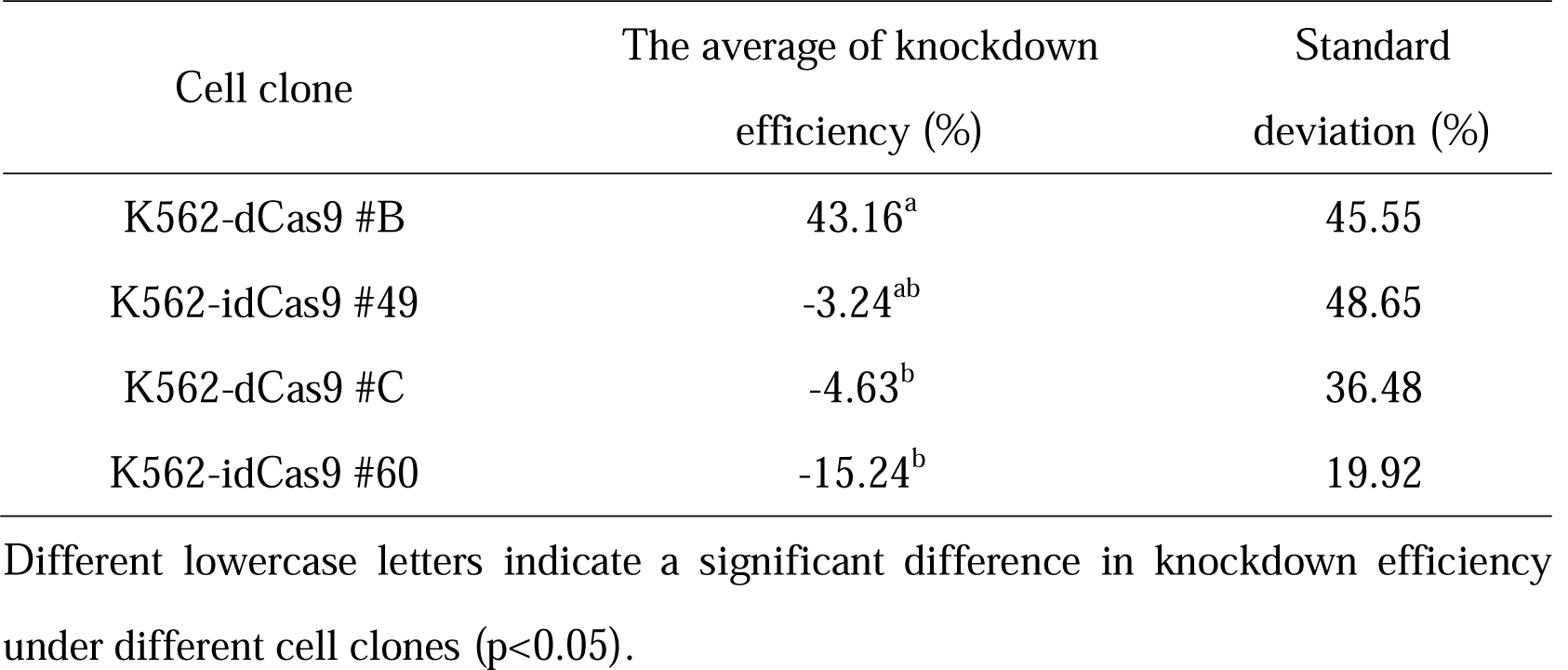
Effects of the four cell clones on knockdown efficiency and multiple comparisons

We also analyzed whether knockdown efficiency is affected by the target gene expression level, and found no correlation (Fig. 3b). The relative expression level of the target genes in four cell clones differed from each other, probably due to the single cloning procedure [11] (Fig. 3c). Continuous monitoring of the KRAB-dCas9 expression revealed that K562-dCas9 #C lost transgene expression over time, while other clones maintained their transgene expression (Supplementary Table 2).

Utilizing K562-dCas9 #B, we assessed the impact of sgRNA expression level on knockdown efficiency. Gene transfer rate, the rate of cells expressing a transgene in the cell population, were assessed to ensure successful transduction. BFP MFI, which is co-expressed with the sgRNA in a polycistronic transcript, was used for determining the gene transfer rate and found to correlate with MOI, fitting a polynomial curve (Fig. 4a). Based on this curve, the gene transfer rate was 34.1% at MOI 0.5, close to the previously reported gene transfer rate 28.6% at MOI 0.5 in K562 cells [12], confirming an effective lentiviral packaging and transduction.

**Fig. 4.**
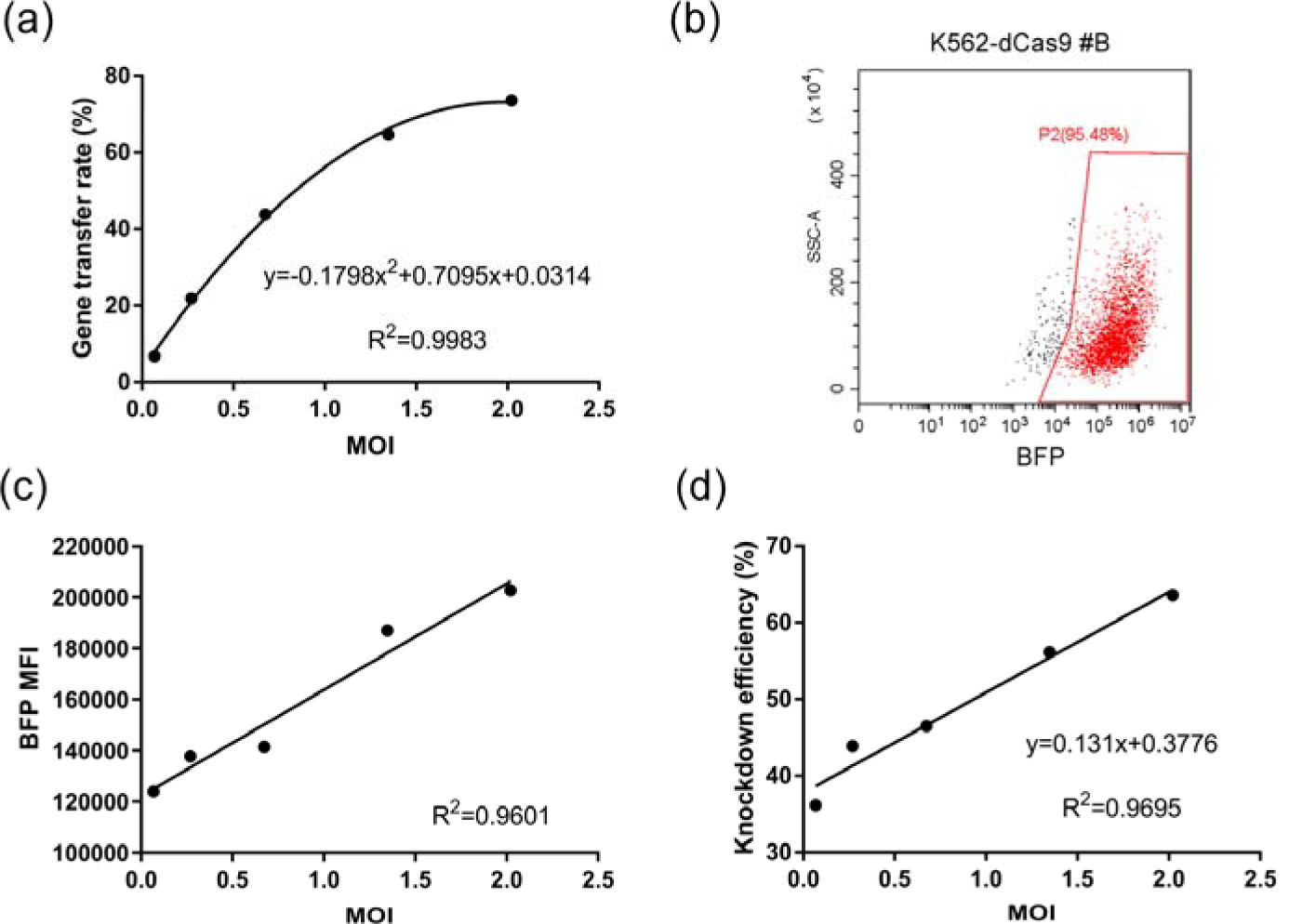
CRISPRi knockdown efficiency correlated with sgRNA expression level. (a) Correlation between MOI of sgRNA infection of the *mmadhc* gene and gene transfer rate, determined by BFP MFI, in K562-dCas9 #B. (b) The scatter plot of K562-dCas9 #B with sgRNA of the *mmadhc* gene. The red frame indicated positive sgRNA expression cells with BFP fluorescence. (c) Correlation between BFP MFI in positive sgRNA expression cells and the MOI of sgRNA infection of the *mmadhc* gene in K562-dCas9 #B. positive sgRNA cells were defined by flow cytometry. (d) Correlation between knockdown efficiency and MOI of sgRNA infection of the *mmadhc* gene in K562-dCas9 #B.

We then assessed how MOI correlated with transgene expression and found that in positive sgRNA expression cells, based on flow cytometry detection(Fig. 4b), BFP MFI linearly correlated with MOI (Fig. 4c). Moreover, a linear relationship was observed between the MOI and the knockdown efficiency (Fig. 4d), indicating that as sgRNA expression level correlates with knockdown efficiency. In our research, based on the linear relationship between the MOI and the knockdown efficiency, around 45% knockdown could be achieved for the *mmadhc* gene by MOI 0.5, the common MOI used for CRISPRi library screening (Fig. 4d).

To systematically evaluate how dCas9 and sgRNA expression level affects the knockdown efficiency, a linear regression model with dCas9 MFI and sgRNA as independent variable individually was applied. Results have shown that the assumption of equal variance was met. When dCas9 serve as the independent variable in the linear model, *R*^*2*^=0.243, *F*=8.367, *p*=0.008, indicating that dCas9 fusion gene expression affected the knockdown efficiency significantly. The standardized coefficient β_1_=0.525 indicated that when dCas9 change for one standard deviation, the knockdown efficiency changes 0.525 standard deviations. Similarly, when sgRNA serves as the independent variable in the linear model, *R*^*2*^=0.949, *F*=75.84, *p*=0.003, and standardized coefficient β_2_=0.981, indicating that sgRNA level affected knockdown efficiency significantly, and when sgRNA level change for one standard deviation, the knockdown efficiency changes for 0.981 standard deviations. The standardized coefficient value has shown that sgRNA level has a greater impact on the knockdown efficiency, which corroborated with the correlation analysis.

## DISCUSSION

As the first step to dissect the dynamics of the KRAB-dCas9 mediated CRISPRi system, we assessed how the expression level of dCas9 fusion protein and the sgRNA affect the knockdown efficiency, which provided the basis for further optimization of the CRISPRi library screening.

We discovered that KRAB-dCas9 expression level significantly impact CRISPRi efficiency, and our data, although limited in the number of clones tested, indicated that enough dCas9 fusion protein probably is a prerequisite for the efficient knockdown. A study on more cell clones is mandated for clarifying whether there is a threshold of KRAB-dCas9 expression above which effective knockdown is guaranteed.

The requirement of the KRAB-dCas9 level has implications for CRISRPi library screenings: its mandated selection of the cells after lentiviral infection, which can be challenging due to many cells required: the number of the cell in each group shall exceed 1000× the sgRNA number in the library. A high-speed cell sorter is required, which is unfortunately not available in most laboratories. CRISPRi library screening without such an instrument might have to rely on obtaining and expanding a single-cell clone that stably expressing a high level of KRAB-dCas9. However, the lentiviral transgene promoter could be methylated and transgene expression shut down in the continuous culture of the cells, as indicated in one of the cell clone K562-dCas9 #C, so continuous monitoring of KRAB-dCas9 expression level is recommended, and more than one selection could be required to obtain such cell clone.

It’s known that single cloning affected the transcription profile of the KRAB-dCas9 expressing cells, and poly-clonal cell populations are better in CRISPRi applications [11]. Effects on transcriptional profile brought by the single cloning procedure can be canceled out, though, as comparisons were made between groups using the same cell clone. Alternatively, with a more potent CRISPRi system, the high expression level of dCas9 fusion protein might not be necessary, and drug selection based on antibiotic resistance could be applied. The dCas9-KRAB-MeCP2 fusion protein is more effective in knockdown [13] and may offer such an option.

Among the six genes randomly chosen for CRISPRi, *npepps* and *rpia* has low efficiency in most or all cell clones. Gene annotation revealed that *npepps* is a puromycin responsive gene, and the puromycin selection after sgRNA transduction probably activated its expression; *rpia* has been reported to be responsive to doxycycline, with increased expression upon doxycycline treatment [14].

We observed that with enough KRAB-dCas9 expression level, CRISPRi efficiency correlated very well with the MOI, which correlated equally well with the sgRNA transgene expression, indicating that CRISPRi efficiency correlated well with sgRNA level. Although it has not been studied in the CRISPRi system, it is known that in the CRISPR system the nuclear concentration of the guide RNA is the limiting factor for efficient DNA targeting [6, 15]. We found that the guide RNA level has a greater impact on CRISPRi efficiency than dCas9. The dynamics of ribonucleoprotein formation and DNA targeting is probably similar between CRISPR and CRISPRi system.

To improve the knockdown efficiency, it is critical to increasing the expression level of sgRNA since sgRNA is very unstable when it was not bound with Cas9 protein [6], improvement of the sgRNA stability could increase efficiency [16]. Alternatively, a change of promotor might serve to increase the sgRNA expression.

The correlation between the sgRNA expression level and the knockdown efficiency of the CRISPRi system has indications for the CRISPRi library screening, too: inducible sgRNA will provide better control of the knockdown. It has been reported that inducible sgRNA offered better control of knockdown in the CRISPR system [17], again pointing to the similarities between the two systems.

## ACKNOWLEDGMENTS

We thank Professor Jonathan S. Weissman, David E. Root and John G. Doench for sharing their CRISPRi library on Addgene, and we acknowledge Dr. Luke A. Gilbert for offering advice on CRISPRi library.

## COMPLIANCE WITH ETHICAL STANDARDS

The authors declare that they have no conflict of interest. This article does not contain any studies involving animals or human participants performed by any of the authors.

