## supplementary for "sgRNA level is a major factor affecting CRISPRi knockdown efficiency in K562 cells"

**Supplementary Table 1.** The sequences of primers

| Primer set | Primer sequence (5’-3’) | Fragment size |
| --- | --- | --- |
| pTRE3G F | tt***gaattc***TTTACTCCCTATCAGTGAT | 392 bp |
| pTRE3G R | ta***gaattcT***TTTACGAGGGTAGGAAGT |  |
| DPH2 qPCR F | ACCTGGACGGAGTGTACGAG | 100 bp |
| DPH2 qPCR R | TCTCCCAATAGCTGGTCAGG |  |
| UBE4A qPCR F | CAGCTGGAAAGCCTCAAATC | 100 bp |
| UBE4A qPCR R | GGCATTTGCAAAAGTGTCCT |  |
| MMADHC qPCR F | CGAACTATCTTCTCCAGCGG | 114 bp |
| MMADHC qPCR R | AAGGCTTTGGGATTGACAACC |  |
| RPIA qPCR F | GCTGGCTATGCTAGTCGCTT | 85 bp |
| RPIA qPCR R | GGATTCCCTTGTGCCACTGA |  |
| ZNF148 qPCR F | TAATGGGTGGAGTGTCTGGC | 85 bp |
| ZNF148 qPCR R | TCCTGGTGAGGCATACTTCG |  |
| LSM4 qPCR F | AGCATCTGTGGACCGTGAA | 86 bp |
| LSM4 qPCR R | GGACCCTGAAGAGAAGGCTA |  |
| NPEPPS qPCR F | TTCTGTGCTGGTGGGTCATA | 127 |
| NPEPPS qPCR R | CATTCATCTCTGGCTTGTCCA |  |
| LIMA1 qPCR F | ATCCCAAGCCACTAAGTCCA | 115 |
| LIMA1 qPCR R | TTCTGACATTCCACGCAGG |  |

Italics indicate restriction endonuclease sites. F: Forward primer, R: Reverse primer.

**Supplementary Table 2. The mCherry MFI in four clones**

|  | K562-dCas9 #B | K562-dCas9 #C | K562-idCas9 #49 | K562-idCas9 #60 |
| --- | --- | --- | --- | --- |
| 1 | 492032.1 | 34448.6 | 21674.1 |  |
| 2 | 389089.4 | 35876.7 | 18261.6 | 761.6 |
| 3 | 419680.5 | 32378.1 | 19004.1 | 1087.9 |
| 4 | 460097.9 | 1779.7 | 19539.1 | 856.5 |

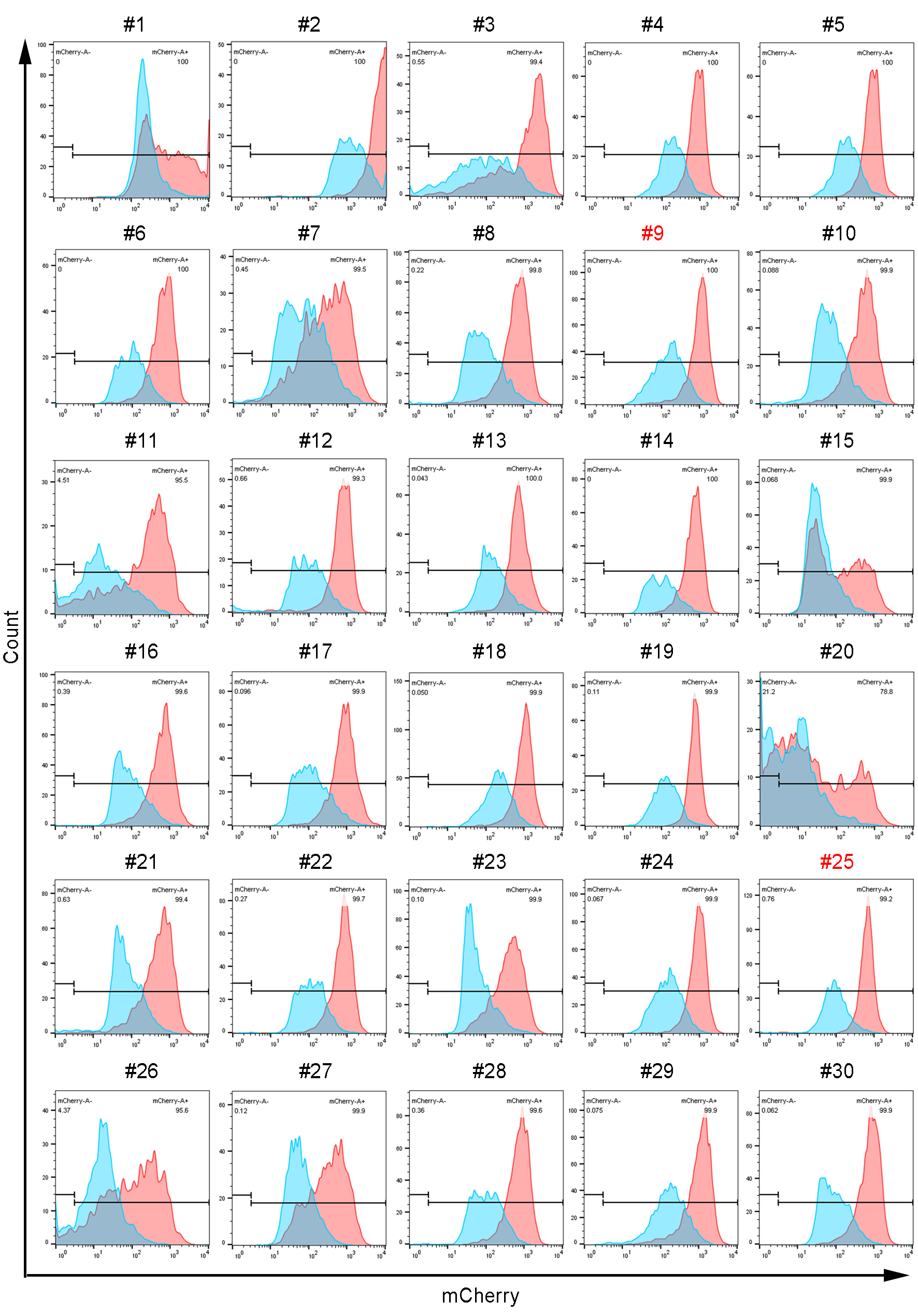

**Supplementary Fig. 1** The histogram of #1 to #30 inducible single clones.

The blue peak is the mCherry fluorescent histogram without induction, and the red peak is with 2000 nM doxycycline induction for 48 h.

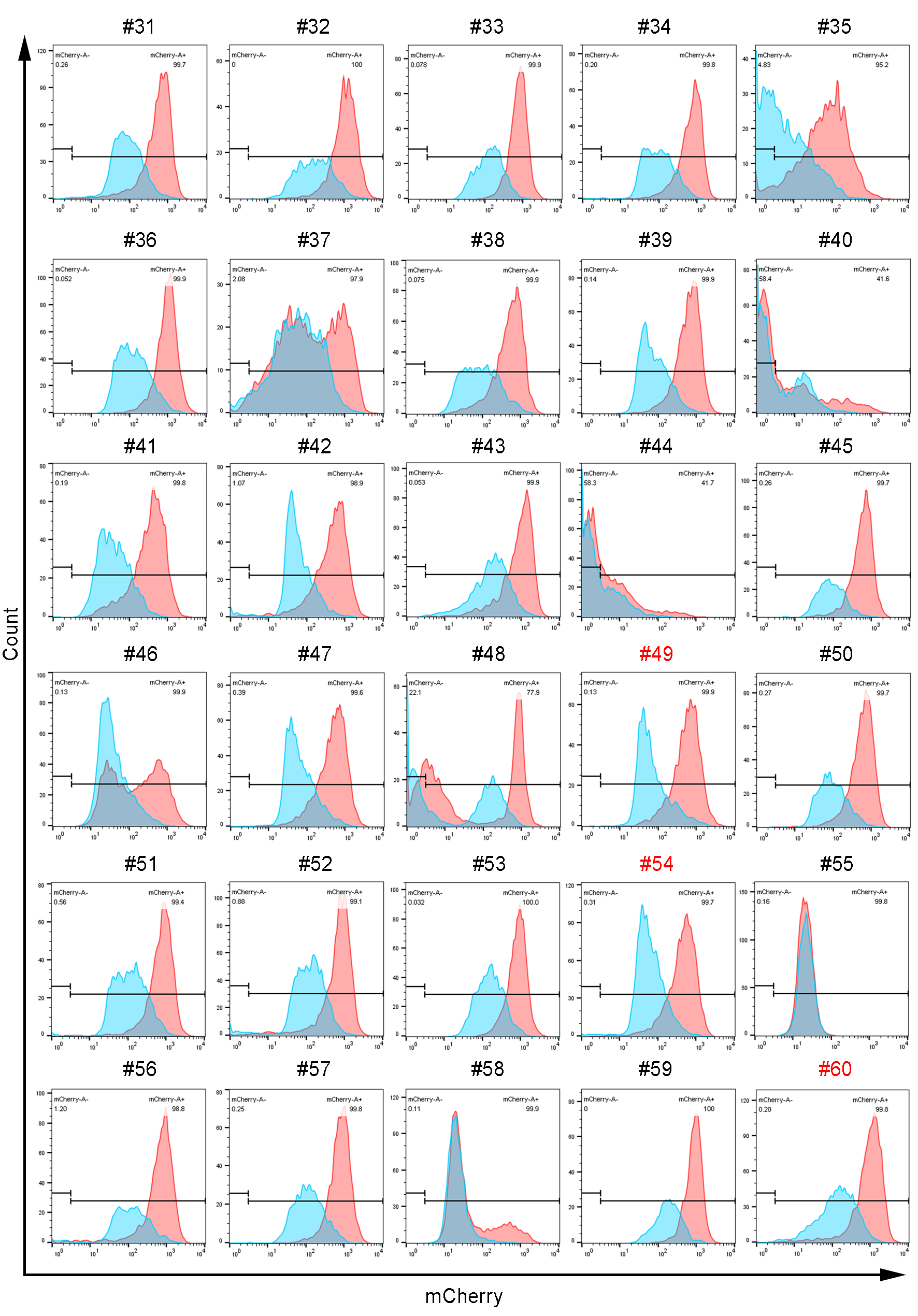

**Supplementary Fig. 2** The histogram of #31 to #60 inducible single clones.

The blue peak is the mCherry fluorescent histogram without induction, and the red peak is with 2000 nM doxycycline induction for 48 h.

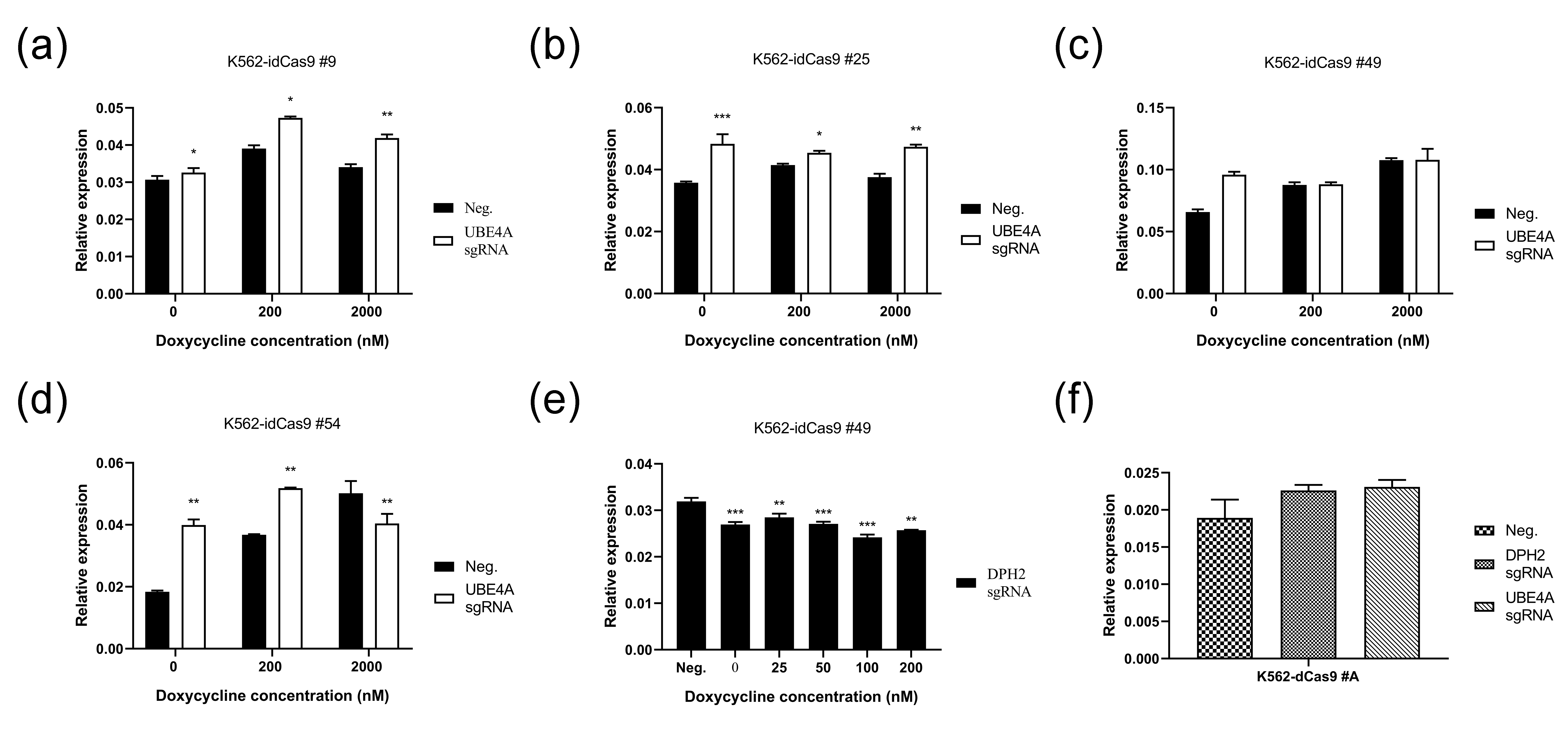

**Supplementary Fig. 3** Pilot CRISPRi experiments targeting UBE4A and DPH2 in cell clones. (a) The relative expression level of UBE4A mRNA in K562-idCas9 #9. (b)The relative expression level of UBE4A mRNA in K562-idCas9 #25. (c) The relative expression level of UBE4A mRNA in K562-idCas9 #49. (d)The relative expression level of UBE4A mRNA in K562-idCas9#54. Neg. indicated none sgRNA transduction. (e) The relative expression level of DPH2 mRNA in K562-idCas9 #49. (f) The relative expression level of UBE4A and DPH2 mRNA in K562-dCas9 #A.
